# Revisiting the genome assembly of *Lupinus* species reveals differential diploidization after a shared whole-genome duplication

**DOI:** 10.64898/2026.04.21.719918

**Authors:** Liying Yang, Yanju Shuai, Weiran Li, Junping Gao, Jianduo Zhang, Haomin Lyu, Songyi Ji, Mingli Chen

## Abstract

Accurate genome assemblies are essential for comparative genomics, yet Hi-C-guided scaffolding can introduce structural errors that misrepresent chromosome architecture and bias evolutionary inferences. Here, we identified pervasive scaffolding errors - including artificial fusions, internal inversions, and incomplete contig mounting - in two previously published *Lupinus* genomes (*L. cosentinii* and *L. digitatus*) using a segmentation method based on LTR retrotransposon density. We re-assembled both genomes, producing chromosome-level references of 472.7 Mb (16 chromosomes) and 427.2 Mb (21 chromosomes), with BUSCO completeness >98.5%. Synteny validation and re-application of LTR profiling confirmed that all prior errors were resolved. Using these corrected genomes together with four additional *Lupinus* species and two outgroup legumes, we investigated post-polyploid evolution. Synonymous substitution rate (Ks) analysis revealed a genus-specific whole-genome duplication (WGD) event (Ks = 0.17) shared by all six *Lupinus* species. The proportion of WGD-derived genes varied markedly, from 60% in *L. digitatus* to only 36% in *L. mutabilis*, indicating differential diploidization. While all species retained a core set of WGD duplicates enriched in cytoskeleton organization, ion transport, and defense responses, each exhibited lineage-specific functional trajectories: cell wall modification in *L. cosentinii* and *L. digitatus*, nitrogen metabolism in *L. albus* and *L. angustifolius*, flower development in *L. luteus*, and stress/lipid metabolism in *L. mutabilis*. Our corrected assemblies provide optimal references for *Lupinus* comparative genomics, and our findings demonstrate that a shared WGD event can lead to both conserved and highly divergent post-polyploid fates, likely underpinning adaptive diversification within the genus.

## Introduction

Recent advances in long-read sequencing and Hi-C technology have enabled chromosome-scale plant genome assemblies (Michael and VanBuren 2020; Kong et al. 2023; Garg et al. 2024; Cheng et al. 2025; Ni et al. 2025). However, Hi-C scaffolding remains prone to structural errors (Burton et al. 2013; Kaul et al. 2020), including artificial chromosome fusions (joining non-syntenic regions), internal inversions (misorientation), and incomplete mounting (missing contigs or gaps). These errors have severe biological consequences: inversions disrupt gene models, fusions create spurious gene clusters, and incomplete mounting leads to missing genes, particularly in repeat-rich regions. Standard completeness metrics like Merqury and BUSCO primarily assess base accuracy and gene content, not chromosomal structural accuracy (Simão et al. 2015; Rhie et al. 2020; Manni et al. 2021), so users may unknowingly draw incorrect conclusions about collinearity, gene family evolution, or adaptive structural variants.

An additional layer of complexity arises from the non-random distribution of transposable elements (TEs) along eukaryotic chromosomes (Wells and Feschotte 2020; Zhang et al. 2020; Viviani et al. 2021). In most plant genomes, long terminal repeat retrotransposons (LTR-RTs) exhibit a characteristic “mountain-shaped” distribution: highly enriched in pericentromeric regions, gradually decreasing towards arms, and lowest near telomeres (Lynch 2007; Venner et al. 2009; Bourque et al. 2018). This predictable pattern serves as an intrinsic quality metric. For instance, in the recently published telomere-to-telomere (T2T) genome of *Lablab purpureus*, prior assemblies showed TE-depleted valleys at putative centromeric positions indicating misassembly, whereas the T2T assembly exhibited TE density peaks precisely at annotated centromeres (Liu et al. 2025). Abnormal TE density profiles—terminal peaks or central troughs—flag scaffolding errors, providing a simple diagnostic tool independent of cross-species synteny.

The genus *Lupinus* (Leguminosae) comprises over 200 species of agronomically and ecologically significant plants, including grain legumes such as white lupin (*L. albus*), narrow-leaf lupin (*L. angustifolius*), and Andean lupin (*L. mutabilis*), as well as wild species adapted to diverse habitats ranging from Mediterranean scrublands to high-altitude Andean plateaus (Sprent et al. 2017; Ishaq et al. 2022; Msaddak et al. 2023). *Lupins* are valued for their high protein content, ability to perform biological nitrogen fixation, and production of unique alkaloids with pharmaceutical potential. In recent years, reference genomes have been published for several *Lupinus* species (Xu et al. 2020; Hufnagel et al. 2021; Pancaldi et al. 2024; Martinez-Hernandez et al. 2025; Susek et al. 2025), but their quality—particularly Hi-C-based assemblies—has not been rigorously evaluated, raising concerns for downstream synteny mapping and inference of structural rearrangements.

A growing body of evidence suggests that the genus *Lupinus* experienced a whole-genome duplication (WGD) event in its common ancestor (Kroc et al. 2014; Cannon et al. 2015; Zhao et al. 2021). Previous studies have proposed a *Lupinus*-specific WGD (Kroc et al. 2014; Susek et al. 2025), but its genomic impact and functional consequences across diverse species have not been systematically characterized using high-quality, error-free genome assemblies. In particular, it remains unknown how post-polyploid gene loss (diploidization) has proceeded in different *Lupinus* lineages, which functional categories are preferentially retained, and whether retention patterns correlate with ecological adaptations (Dodsworth et al. 2016; Mandáková and Lysak 2018; Qiao et al. 2019).

In this study, we developed a segmentation method exploiting LTR retrotransposon mountain-shaped distribution to detect scaffolding errors. Applying this method to two previously published *Lupinus* genomes (*L. cosentinii* and *L. digitatus*) (Susek et al. 2025), we identified multiple problematic scaffolds. Validation and re-assembly using the same raw PacBio HiFi and Hi-C data confirmed errors originated from Hi-C scaffolding. We generated de novo chromosome-level assemblies for both species (N50 > 10 Mb, BUSCO > 98.5%). Using these corrected genomes as references, we then performed a genus-wide comparative analysis to investigate the evolutionary consequences of polyploidy.

## Materials and Methods

### 1. Error detections of genome assembly for two *Lupinus* genomes

The genomes of *Lupinus cosentinii* and *L. digitatus* were obtained from previously published literature (Susek et al. 2025). Then, LTR retrotransposons were annotated using the LTR_FINDER combined with LTR_retriever pipeline (Xu and Wang 2007; Ou and Jiang 2018). Custom scripts were then used to process the annotation information. Each chromosome was divided into 100 windows (window size: 3 Mb), and the density of LTR retrotransposons was calculated per window. The TE density sequence of each chromosome was equally partitioned into four segments, and the ratio of the average LTR density at the two terminal segments to the average density in the middle two segments was computed. This ratio, together with visualizations of LTR distributions, was used to judge the assembly correctness of each chromosome. Chromosomes showing abnormal segmental LTR densities or distributions were considered problematic.

### 2. Genome assembly and chromosome-scale scaffolding

Genome assembly for *L. cosentinii* and *L. digitatus* was performed using a strategy comprising initial contig assembly and chromosome anchoring via Hi-C data. The Pacbio HiFi and Hi-C data for the two species were obtained from the NCBI repository under the BioProject number PRJNA1080360 (Susek et al. 2025). Contig assembly was carried out using hifiasm (v0.25.0-r726) with default parameters (Cheng et al. 2026 Feb 4). Hi-C reads were mapped to the primary contigs using Chromap (v0.2.6-r490) (Zhang et al. 2021), and chromosome construction was performed with YaHS (v1.2a.1) (Zhou et al. 2023). A contact matrix was generated using Juicer (v1.1), followed by manual curation in Juicebox (v1.11.08) to rectify misassemblies and refine scaffold orientations (Durand et al. 2016; Robinson et al. 2018). Genome completeness was assessed using BUSCO v5.8.3 with the embryophyta_odb12 and eudicotyledons_odb12 lineage datasets (Simão et al. 2015; Manni et al. 2021).

### 3. Annotations of the two re-assembled genomes

Repetitive sequence annotation in both *L. cosentinii* and *L. digitatus* genomes was conducted using the EarlGrey pipeline under default parameters (Baril et al. 2024). Prior to gene calling, repetitive regions were masked with RepeatMasker (v4.1.5) (http://www.repeatmasker.org). A three-pronged annotation strategy was adopted, integrating homology-based searches, transcriptome-supported evidence, and ab initio algorithms. For homology-based inference, protein sequences from *Arabidopsis thaliana* (Araport11) and *Glycine max* (Wm82) were retrieved from Phytozome (Cheng et al. 2017; Espina et al. 2024), and gene models were generated using GeMoMa (v1.9) (Keilwagen et al. 2019). Transcriptome evidence incorporated short-read RNA-seq datasets from PRJNA1080360, aligned with Hisat2 (v2.2.1) (Kim et al. 2019), followed by transcript isoform reconstruction with TACO (v0.7.3) (Niknafs et al. 2017) and coding sequence extraction with TransDecoder (v5.7.0) (https://github.com/TransDecoder/TransDecoder). *De novo* prediction was executed with Braker2 (v2.1.2) (Brůna et al. 2021), integrating Augustus (v3.3.2) (Stanke et al. 2006) and GeneMark-ET (v4.59) (Brůna et al. 2024), trained using RNA-seq alignments and Swiss-Prot protein sequences. Final gene sets were derived by merging the three prediction streams through EVidenceModeler (v2.0.0) (Haas et al. 2008). Functional characterization of predicted genes, including GO and KEGG categories, was performed using eggNOG-mapper (v2.1.12) (Cantalapiedra et al. 2021) against the eggNOG database (v5.0.2) (Huerta-Cepas et al. 2018).

### 4. Verification of re-assembled *Lupinus* genomes

The accuracy of the Hi-C-guided scaffolding was assessed through whole-genome synteny comparison between the newly assembled genomes and their previously published versions using D-Genies. Scaffolding errors were categorized into chromosome fusions, internal inversions (orientation errors), and incomplete contig mounting (fragmented or missing alignments). Additionally, the LTR density-based segmentation method described as above was reapplied to the re-assembled genomes to confirm that all previously flagged problematic scaffolds had been correctly resolved.

### 5. Phylogeny analysis of *Lupinus* genus

Protein sequences from *L. cosentinii, L. digitatus*, and six additional legume species (*Ammopiptanthus mongolicus, Crotalaria pallida, L. albus, L. angustifolius, L. luteus*, and *L. mutabilis*) (Xu et al. 2020; Hufnagel et al. 2021; Feng et al. 2024; Pancaldi et al. 2024; Martinez-Hernandez et al. 2025; Yang et al. 2026) were subjected to orthologous clustering using OrthoFinder (v2.4) (Emms and Kelly 2019) with DIAMOND alignment (Buchfink et al. 2021). A maximum-likelihood species tree was constructed based on single-copy orthologous genes shared across all ten species. Multiple sequence alignments were generated with MAFFT v7.205 (parameters: ‘--localpair --maxiterate 1000’) (Katoh and Standley 2013). IQ-TREE v1.6.11 was then employed for tree inference using default parameters (Nguyen et al. 2015). Branch support was evaluated using 1,000 bootstrap replicates. Divergence times were estimated with the MCMCTree program in the PAML package (Yang 2007), using *A. mongolicus* as the outgroup. The calibration timepoints were obtained from the TimeTree database (Kumar et al. 2022).

### 6. Gene family expansion and contraction

Dynamics of gene family size were analyzed using CAFE v4.2 (Mendes et al. 2021). The software was provided with the phylogenetic tree topology and the gene family counts derived from MCMCTree and OrthoFinder. A birth-death model was fitted to infer ancestral gene family sizes, and significant expansion or contraction events along each modern species branch were identified using a likelihood ratio test (p < 0.05).

### 7. Whole-genome duplication (WGD) analysis

Paleopolyploidy events were investigated through comparative synteny analysis and synonymous substitution rate (Ks) modeling. Syntenic blocks with the eight genomes (*L. cosentinii, L. digitatus, Ammopiptanthus mongolicus, Crotalaria pallida, L. albus, L. angustifolius, L. luteus*, and *L. mutabilis*) were identified using MCScanX (Wang et al. 2012). Homologous gene pairs within collinear blocks were extracted, and their Ks values were calculated using the KaKs_Calculator 2.0 (Wang et al. 2010). Ks distribution histograms were plotted to detect peaks corresponding to ancient WGD events.

### 8. Post-WGD divergence of the gene space

Based on the syntenic analyses described above, WGD-derived (paralogous) genes were identified within each of the six *Lupinus* genomes (*L. cosentinii, L. digitatus, L. albus, L. angustifolius, L. luteus*, and *L. mutabilis*). We then examined the distribution of these WGD-derived genes among gene families that showed significant expansion. For each species, the proportion of expanded-family genes originating from WGD was calculated.

To explore functional divergence following polyploidy, we performed Gene Ontology (GO) and KEGG pathway enrichment analyses using R package “clusterprofiler” (Guangchuang Yu. et al. 2012), specifically on the WGD-derived genes within expanded families across the six species. Enrichment patterns revealed both conserved and lineage-specific functional trajectories following the WGD event in the common ancestor of *Lupinus* genus.

## Results

### 1. Detection of assembly errors in *Lupinus cosentinii* and *L. digitatus*

To identify potential assembly errors, we applied a segmentation-based scoring method using the chromosomal distribution of LTR retrotransposons. Suspected regions were further validated through manual inspection of LTR distribution plots (Figure 1), re-assembly validation using raw sequencing reads, and synteny comparison with closely related reference genomes (Figure 3).

**Figure 1.**
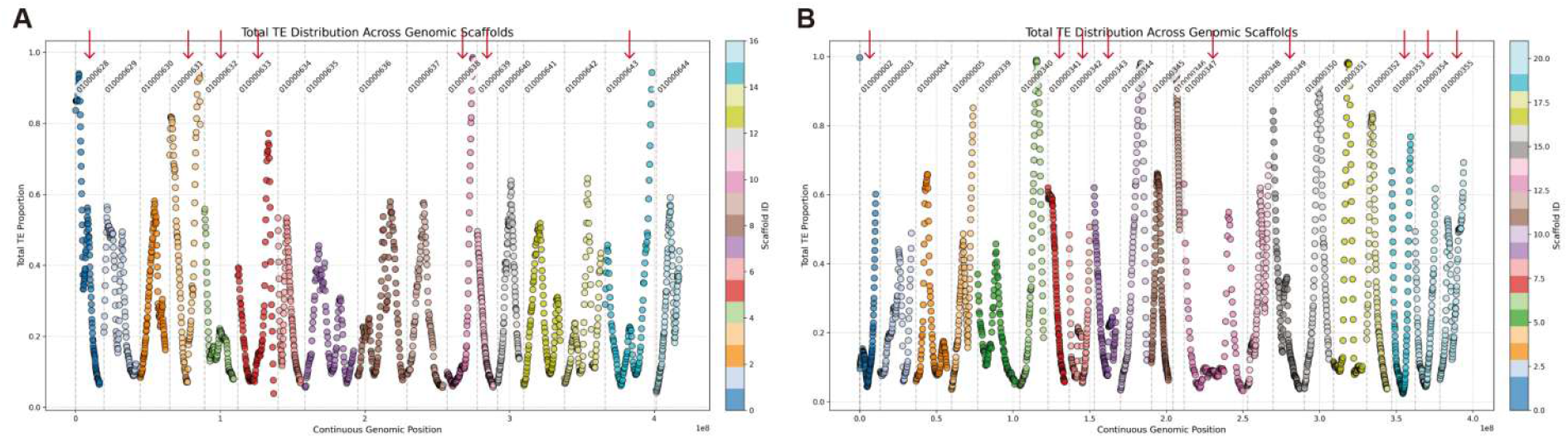
Identification of scaffolding errors in *Lupinus cosentinii* and *L. digitatus* using transposon distributions. (a) LTR density profiles for seven problematic scaffolds of *L. cosentinii* (scaffolds 628, 631, 632, 633, 638, 639, 643; full identifiers e.g., JBANGR010000631.1). (b) LTR density profiles for nine problematic scaffolds of *L. digitatus* (scaffolds 2, 341, 342, 343, 347, 349, 353, 354, 355; e.g., JBANGQ010000353.1).

In the previously published *L. cosentinii* genome, seven scaffolds were detected as problematic: 628, 631, 632, 633, 638, 639, and 643 (full identifiers e.g., JBANGR010000631.1; Figure 1A). For *L. digitatus*, nine scaffolds were flagged: 2, 341, 342, 343, 347, 349, 353, 354, and 355 (e.g., JBANGQ010000353.1; Figure 1B). Notably, the actual number of assembly errors exceeded these detected cases, indicating that our approach, while effective, does not capture all structural misassemblies.

Further inspection revealed that most errors originated from the Hi-C-guided scaffolding step. Incorrect joins between non-syntenic regions, misoriented contigs, and spurious fusions of telomeric or centromeric repeats led to erroneous chromosome-level mounting (Figure 1). Such scaffolding mistakes can severely compromise downstream applications, including gene order analysis, synteny-based comparative genomics, and accurate interpretation of chromosomal rearrangements. Users of these genome assemblies should therefore exercise caution when drawing conclusions about collinearity or structural variants involving the flagged scaffolds.

### 2. Genome re-assemblies of the *L. cosentinii* and *L. digitatus*

The genomes of *L. cosentinii* and *L. digitatus* were re-assembled using PacBio HiFi reads and Hi-C data (BioProject PRJNA1080360), followed by chromosome-level scaffolding as described in Materials and Methods. For *L. cosentinii*, the initial contig assembly consisted of 871 contigs with an N50 of 12.77 Mb, indicating high continuity prior to scaffolding (Figure 2A; Supplementary Table 1). The final assembly comprised 16 chromosomes with a total length of 472.7 Mb. Individual chromosome sizes ranged from 18.03 Mb (chr12) to 38.91 Mb (chr09). For *L. digitatus*, 21 chromosomes were obtained, covering 427.2 Mb, with sizes ranging from 14.22 Mb (chr15) to 39.91 Mb (chr03) (Figure 2B; Supplementary Table 1). The underlying contig set contained 580 sequences (N50 = 10.05 Mb), reflecting similarly high assembly contiguity.

**Figure 2.**
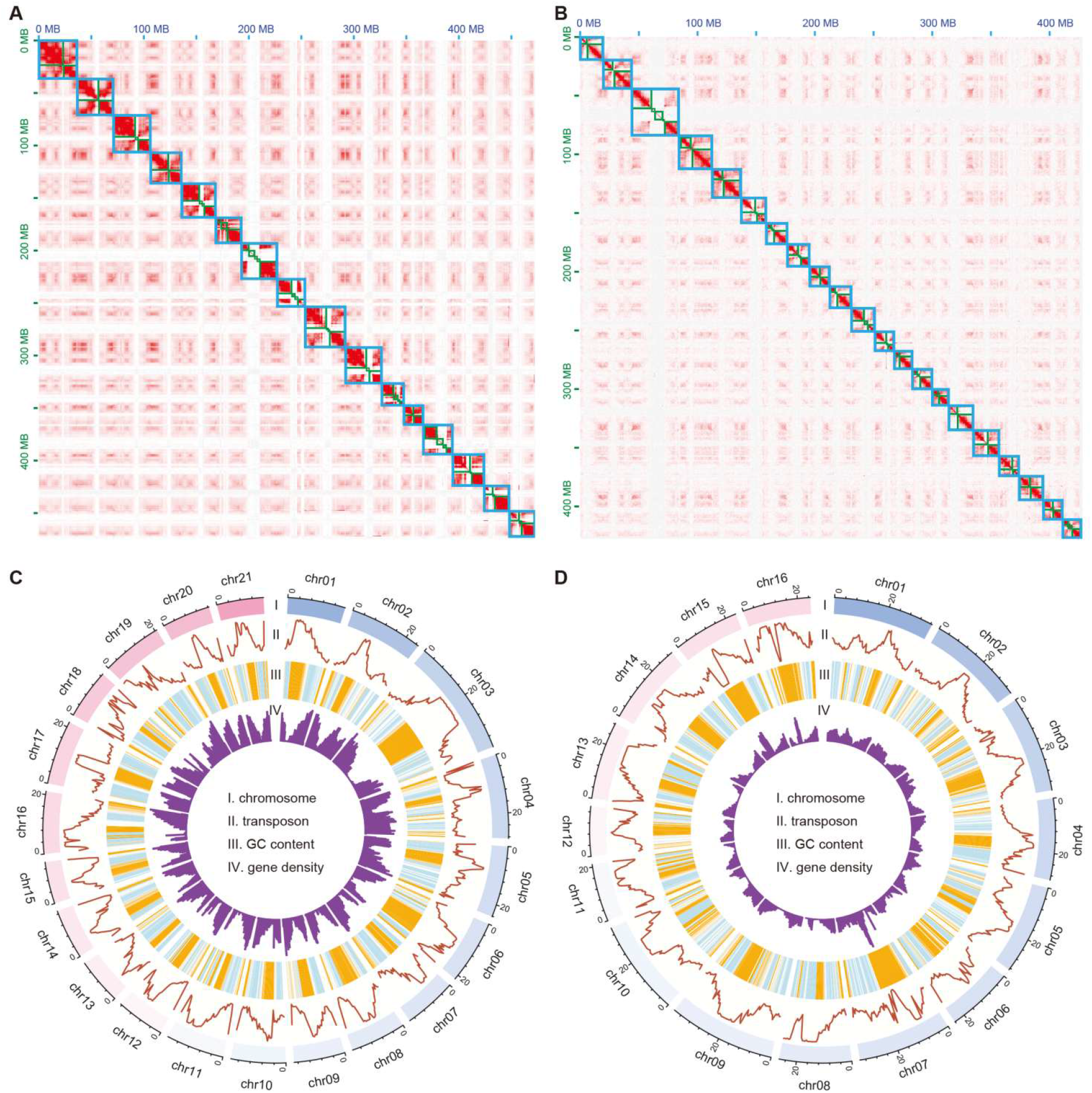
Chromosome-scale features of the reassembled *Lupinus cosentinii* and *L. digitatus* genomes. (a) Heatmap of Hi-C interaction for the *L. cosentinii* genome and scaffolding. (b) Heatmap of Hi-C interaction for the *L. digitatus* genome and scaffolding. (c) Circos plot of the re-assembled *L. cosentinii* genome. (d) Circos plot of the re-assembled *L. digitatus* genome.

Genome completeness was assessed using BUSCO v5.8.3 with two plant-specific lineage datasets. *L. cosentinii* showed 98.6% complete BUSCOs (embryophyta_odb12: 77.9% single-copy, 20.7% duplicated) and 98.5% complete BUSCOs (eudicotyledons_odb12: 74.5% single-copy, 24.0% duplicated) (Supplementary Table 2). *L. digitatus* achieved 98.8% (embryophyta: 79.3% single-copy, 19.5% duplicated) and 98.5% (eudicots: 75.5% single-copy, 23.0% duplicated) (Supplementary Table 2). The high duplication rates in both genomes (24.0% in *L. cosentinii* and 23.0% in *L. digitatus* for eudicots) suggest substantial retention of duplicated genes.

### 3. Genome annotation and repeat composition

Protein-coding gene annotation integrated homology-based, transcriptome-supported and *ab initio* predictions. A total of 36,545 genes were annotated in *L. cosentinii* and 32,972 in *L. digitatus* (Figure 2C-D; Supplementary Table 3). Functional annotation using eggNOG-mapper assigned putative functions to 87.1% and 92.7% of the predicted genes, respectively (Figure 2C-D; Supplementary Table 3).

In *L. cosentinii*, interspersed repeats covered 52.80% of the genome (249.6 Mb). Retroelements were the dominant class (23.39%), with LTR retrotransposons accounting for 21.73% (Ty1/Copia: 13.92%; Gypsy/DIRS1: 7.72%) (Figure 2C; Supplementary Table 4). LINEs represented 1.66%, while DNA transposons contributed 1.87%. By contrast, *L. digitatus* showed a lower total interspersed repeat content (41.74%; 178.3 Mb) (Figure 2D; Supplementary Table 5). Retroelements comprised 17.24% (LTRs: 15.53%; Ty1/Copia: 9.25%; Gypsy: 6.13%), whereas DNA transposons were more abundant (3.26%), driven primarily by MULE-MuDR elements (2.06%). The most striking difference was satellite DNA: *L. digitatus* contained 10.79% satellites (46.1 Mb), compared to only 0.41% in *L. cosentinii*. This large variation may reflect differential centromere evolution or repeat expansion dynamics following speciation. The high-quality assemblies and annotations presented here provide a robust foundation for comparative genomic analyses within the genus *Lupinus*.

### 4. Synteny validation of the reassembled genomes

To assess the accuracy of our Hi-C-guided scaffolding and to identify residual assembly errors, we compared the newly assembled genomes of *L. cosentinii* and *L. digitatus* with their previously published versions using D-Genies (Figure 3). Errors identified in the previous assemblies originated from the Hi-C scaffolding step despite manual curation in Juicebox and could be classified into three categories: chromosome fusions (artificially merging non-syntenic regions), orientation errors (internal inversions affecting gene orientation and regulatory annotation), and incomplete mounting (missing genes or broken chromosomes). All observations were supported by Hi-C interaction heatmaps (Figure 2A-B).

**Figure 3.**
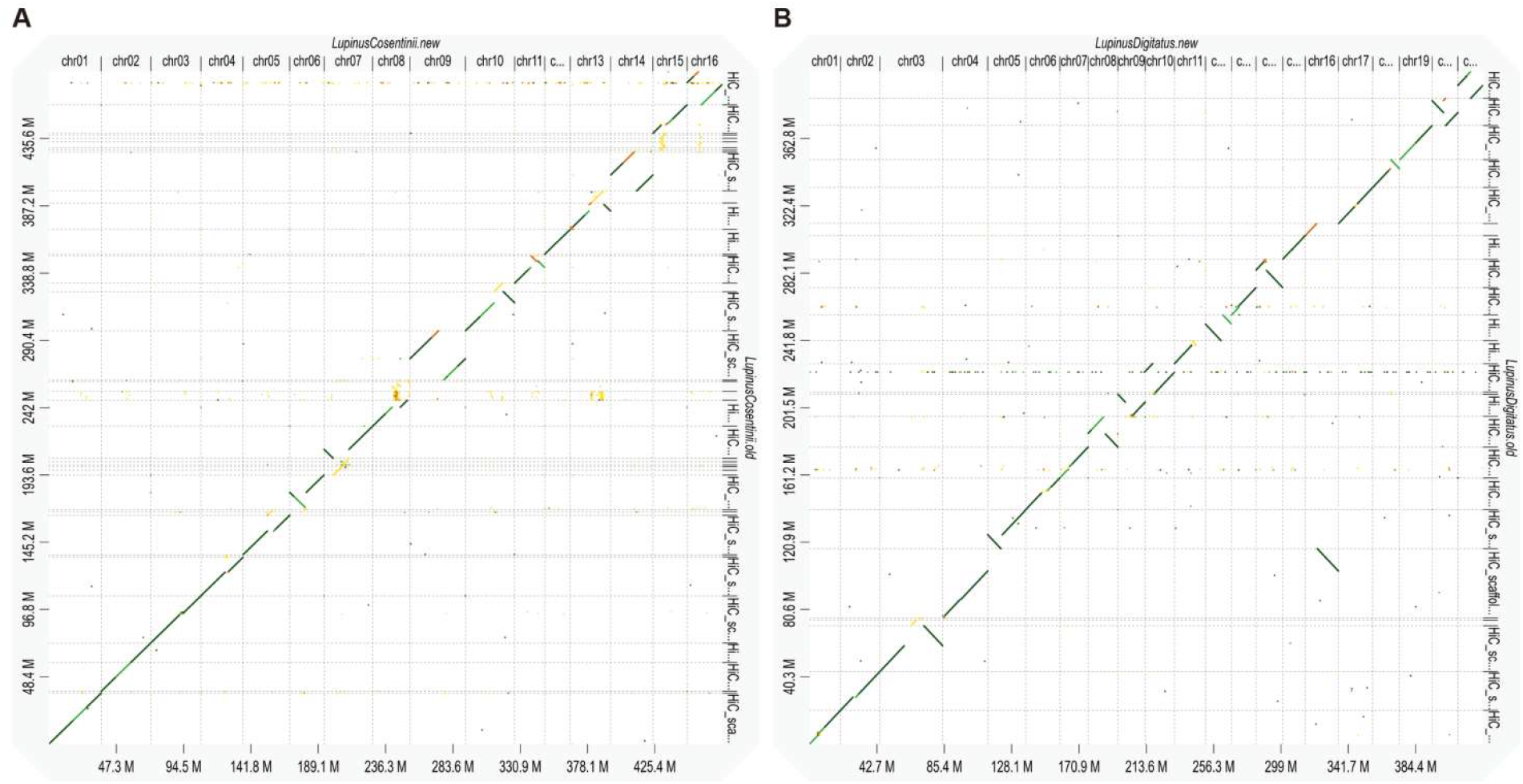
Whole-genome synteny comparison between the re-assembled genomes and the previously published assemblies for *Lupinus cosentinii* (a) and *L. digitatus* (b).

Synteny analysis for *L. cosentinii* revealed that out of 16 chromosomes in the new assembly, 11 exhibited scaffolding errors in the previous version, while the remaining 5 were correctly assembled (Figure 3A). Eight chromosomes (chr06, chr07, chr09, chr10, chr11, chr13, chr14, and chr16) displayed a continuous diagonal but with internal segments reversed (anti-diagonal). For example, our newly assembled chr09 corrected the orientations of both arms, which was supported by the Hi-C interaction heatmap. Incomplete contig mounting was observed in six chromosomes (chr05, chr07, chr08, chr10, chr13, and chr15).

For *L. digitatus*, the new assembly contains 21 chromosomes, of which synteny comparison identified 12 chromosomes with assembly errors (Figure 3B). Artificial chromosome fusions has happened to chr04 and chr16. Internal inversions, representing the orientation errors, were observed in 11 chromosomes, including chr03, chr05, chr08, chr09, chr10, chr11, chr12, chr14, chr18, chr20, and chr21. Incomplete contig mounting could be found in chr03 and chr11, which had alignment gaps or highly fragmented diagonals, confirming that some contigs were left unplaced.

### 5. Validation of corrected assembly errors using LTR distribution analysis

To verify that the scaffolding errors identified in the previously published genomes (Figure 1) have been successfully resolved in our re-assembled versions, we applied the same segmentation-based scoring method to evaluate LTR retrotransposon distribution across all chromosomes (Figure 4).

**Figure 4.**
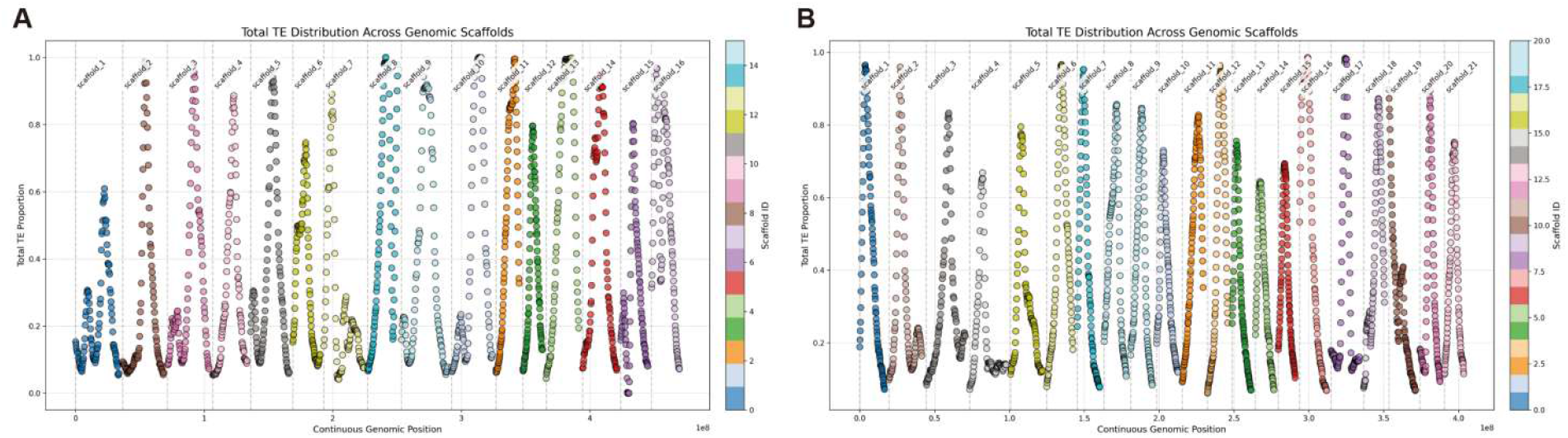
Validation of corrected assembly errors using transposon density profiles in the reassembled genomes of *Lupinus cosentinii* (a) and *L. digitatus* (b).

For *L. cosentinii*, all 16 chromosomes exhibited a normal mountain-shaped LTR density pattern, with decreased repeat densities at the terminal regions and elevated densities in the central portions (Figure 4A). No chromosome was flagged as abnormal by our scoring method. Specifically, the previously problematic scaffolds (628, 631, 632, 633, 638, 639, 643) have been correctly incorporated into their respective chromosomes without any remaining fusion, inversion, or incomplete mounting errors. Similarly, for *L. digitatus*, all 21 chromosomes displayed clean, symmetric LTR distribution profiles (Figure 4B). The nine previously problematic scaffolds (2, 341, 342, 343, 347, 349, 353, 354, 355) were no longer detectable as independent entities; instead, they now reside within correctly oriented and fully mounted chromosomes. Importantly, additional assembly errors that were not captured by the original detection pipeline (e.g., subtle misjoins or missing contigs in the old assemblies) have also been rectified. These results confirm that our re-assembly and manual curation efforts have produced high-quality, error-resolved genome references for both *Lupinus* species.

### 6. Whole-genome duplication and retention of WGD-derived genes

Synteny-based synonymous substitution rate (Ks) analysis was performed on collinear gene pairs from six *Lupinus* species and two outgroup legumes (*Ammopiptanthus mongolicus, Crotalaria pallida*). All six *Lupinus* genomes shared a prominent Ks peak at ∼0.17, absent in outgroups (Figure 5A), indicating a genus-specific whole-genome duplication (WGD) event in the common ancestor of *Lupinus*. This is supported by BUSCO assessments: all Lupinus genomes contained >20% duplicated genes, versus <8% in outgroups (Supplementary Table 2).

**Figure 5.**
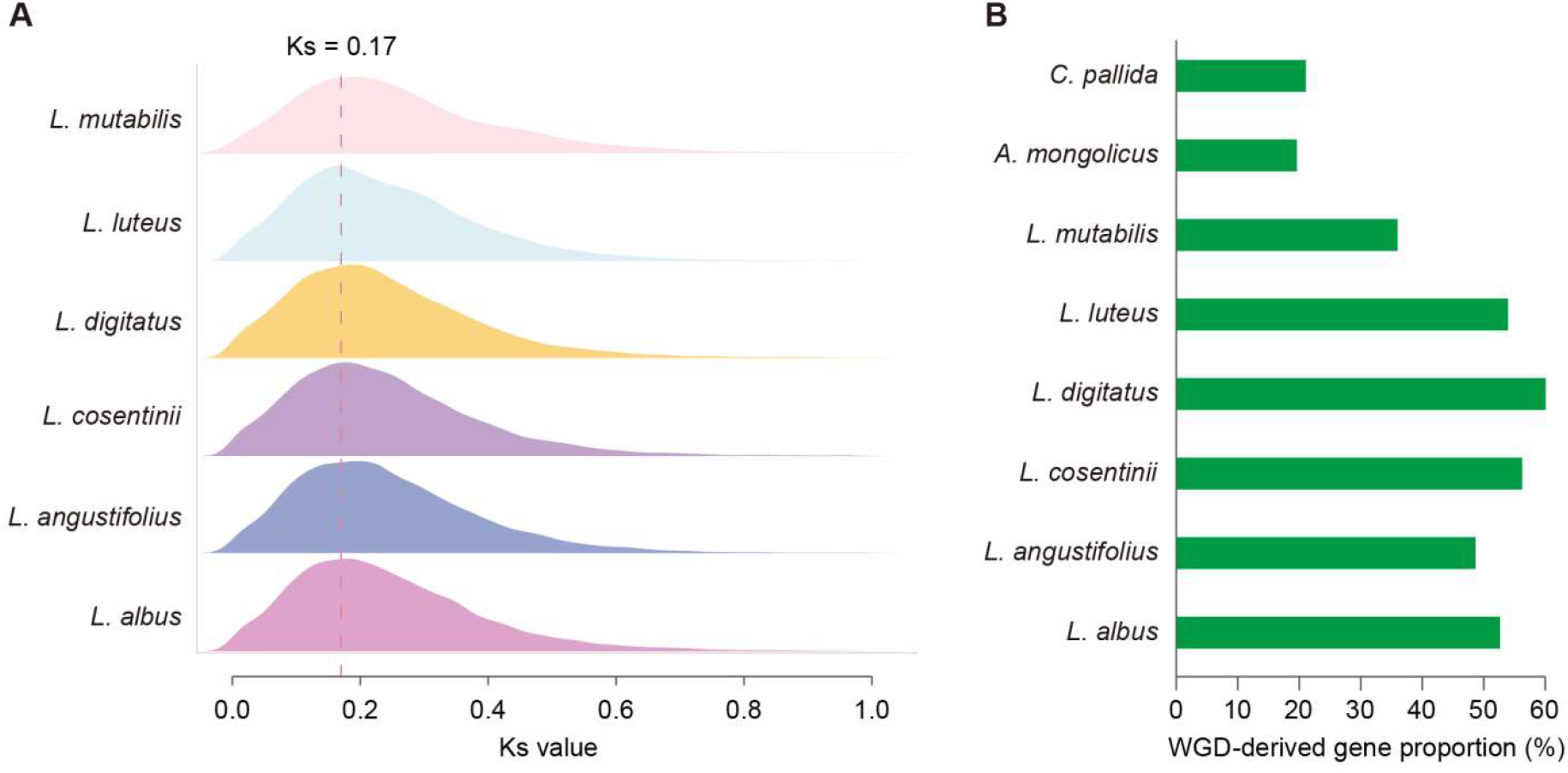
Whole-genome duplication (WGD) event in *Lupinus* and the proportion of WGD-derived genes across eight legume species. (a) Ks (synonymous substitution rate) distribution of syntenic gene pairs within each genome. A prominent peak at Ks = 0.17 is observed in all six *Lupinus* species. (b) Proportion of WGD-derived genes (percentage of total protein-coding genes) in each species.

The proportion of WGD-derived paralogs varied substantially among species (Figure 5B; Supplementary Table 6). *L. digitatus* showed the highest retention (60.08%, 19,810/32,972 total genes), followed by *L. cosentinii* (56.25%, 20,557/36,545), *L. luteus* (53.95%, 19,900/36,884), *L. albus* (52.64%, 20,139/38,255), and *L. angustifolius* (48.66%, 16,094/33,074). In contrast, *L. mutabilis* retained only 35.99% (13,586/37,754). Outgroup values were 19.62% and 21.08%. Thus, the *Lupinus*-specific WGD contributed a large fraction of gene repertoires in most species, with retention exceeding 50% in five of six species, highlighting substantial polyploidy-driven gene family expansion.

### 7. Biased retention and functional divergence of WGD-derived genes

The *Lupinus* -specific WGD (Ks = 0.17) maps to the branch leading to the common ancestor of all six *Lupinus* species (red star in Figure 6A). Across all eight species, contracted gene families consistently outnumber expanded families (Figure 6B; Supplementary Table 6), indicating widespread post-WGD gene loss (diploidization). This pattern is especially pronounced in *Lupinus*, where contractions (336–931) exceed expansions (112–264) by 1.5- to 3-fold. Species-specific variation (e.g., *L. mutabilis*: 667 contractions vs. 112 expansions) points to lineage-specific diploidization trajectories.

**Figure 6.**
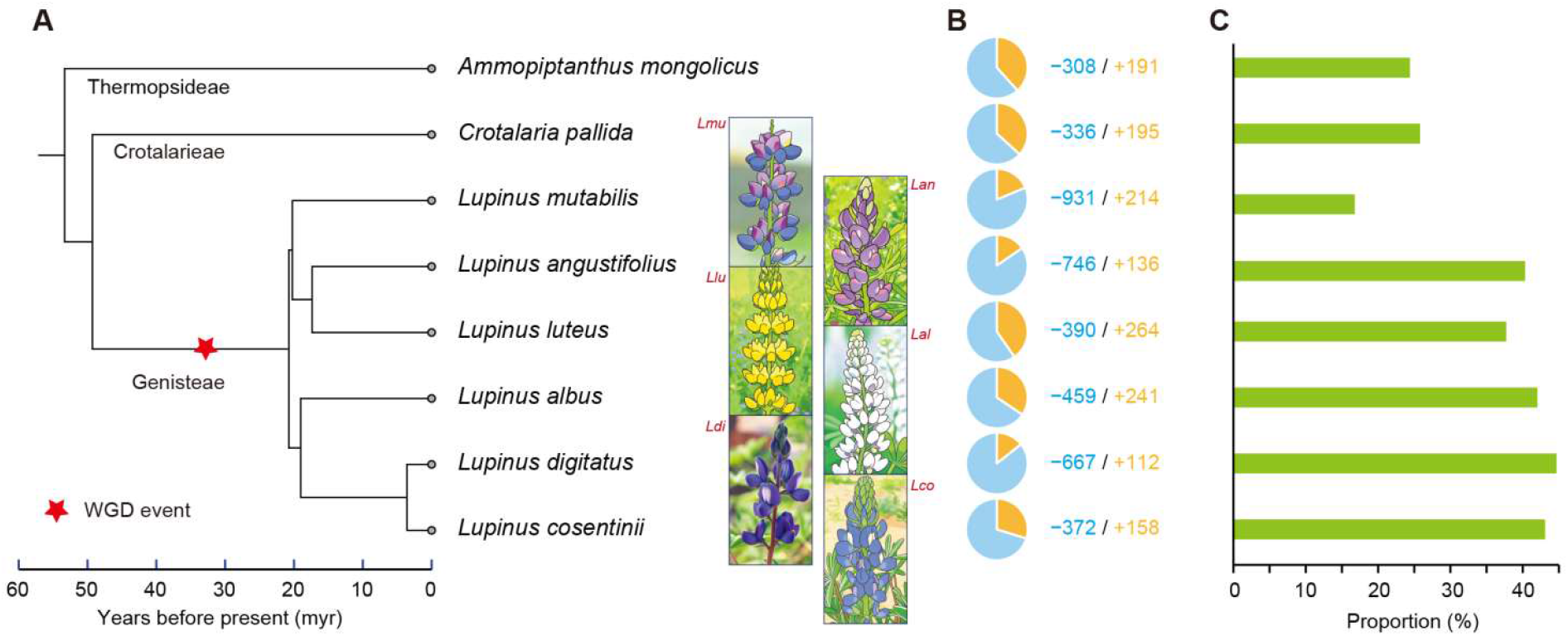
Phylogeny, gene family dynamics, and contribution of WGD to expanded families across eight legume species. (a) Phylogenetic tree and divergent age across eight genomes. The red star marks the genus-specific whole-genome duplication (WGD) event (Ks = 0.17). (b) Numbers of significantly expanded (red) and contracted (blue) gene families per species (likelihood ratio test, p < 0.05). The values indicate the exact counts for significant contraction and expansion families (e.g., −308/+191 for *A. mongolicus*). (c) Proportion of WGD-derived genes within significantly expanded gene families.

The contribution of WGD-derived genes to significantly expanded families also varies (Figure 6C; Supplementary Table 6). *L. digitatus* (44.67%), *L. cosentinii* (43.10%), *L. albus* (42.03%), and *L. angustifolius* (40.33%) show high proportions, *L. luteus* intermediate (37.72%), while *L. mutabilis* exhibits a strikingly low value (16.78%), comparable to outgroups (24.40–25.80%). GO enrichment analysis of WGD-derived genes within expanded families reveals a conserved core across all six species: actin cytoskeleton organization, ion transport, calcium signaling, and defense responses (Figure 7). However, each species displays unique signatures. *L. cosentinii* and *L. digitatus* additionally enrich for cell wall modification and root hair development (Figure 7A-B; Supplementary Tables 7-8). *L. albus* and *L. angustifolius* show distinct enrichment in nitrogen compound metabolism and symbiotic nitrogen fixation (Figure 7C-D; Supplementary Tables 9-10). *L. luteus* uniquely enriches flower development and seed regulation terms (Figure 7E; Supplementary Table 11). Strikingly, *L. mutabilis* lacks the universal cytoskeleton/ion transport/calcium enrichments; instead, its GO terms are dominated by MAPK cascade, fatty acid/vitamin E biosynthesis, oxidative stress, and protein processing (Figure 7F; Supplementary Table 12). These results demonstrate that after the shared WGD, most species retained a core set of functions, but each followed lineage-specific trajectories—with *L. mutabilis* representing an extreme case of post-polyploid functional divergence.

**Figure 7.**
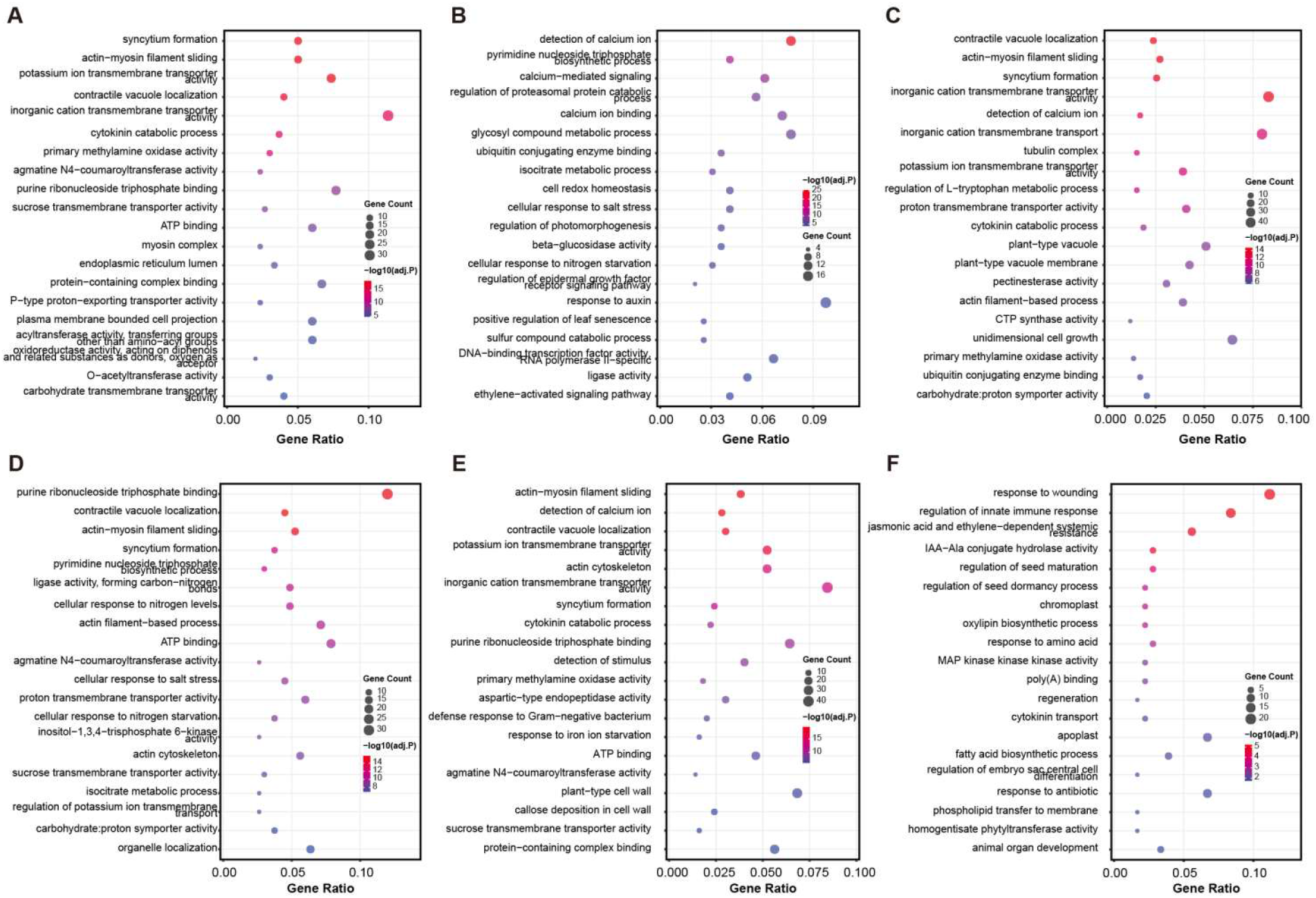
Functional divergences of the WGD-derived genes within significantly expanded gene families. Only significantly enriched GO categories were presented for each genome (p.adjust < 0.05). The results of GO enrichment presented for the six *Lupinus* genomes: (a) *L. cosentinii*, (b) *L. digitatus*, (c) *L. albus*, (d) *L. angustifolius*, (e) *L. luteus*, (f) *L. mutabilis*.

## Discussion

Accurate genome assemblies are fundamental to comparative genomics. Here, we identified pervasive scaffolding errors in two published *Lupinus* genomes (*L. cosentinii, L. digitatus*), re-assembled using both PacBio HiFi and Hi-C data, and conducted a genus-wide analysis of post-polyploid evolution. Our results reveal the vulnerability of Hi-C-guided scaffolding, provide high-quality references for *Lupinus*, and demonstrate both conserved and lineage-specific patterns following a shared whole-genome duplication (WGD).

Using an LTR retrotransposon density-based segmentation method, we detected three error types in the original assemblies: artificial chromosome fusions, internal inversions, and incomplete contig mounting. Despite manual curation in Juicebox, most errors originated from Hi-C scaffolding. Such mistakes have biological consequences: inversions disrupt gene models and syntenic relationships; fusions create spurious gene clusters; incomplete mounting leads to missing genes, especially in repeat-rich regions. Our findings echo growing concerns that Hi-C scaffolding requires rigorous validation. The LTR-based scoring method offers a simple diagnostic tool for plant genomes, where LTR retrotransposons are abundant and typically exhibit mountain-shaped chromosomal distributions.

We generated *de novo* assemblies for *L. cosentinii* (16 chromosomes, 472.7 Mb) and *L. digitatus* (21 chromosomes, 427.2 Mb) with improved parameters and intensive curation. Synteny comparisons and re-application of LTR profiling confirmed that all previously flagged errors were resolved. Both assemblies show high contiguity (N50 > 10 Mb) and completeness (BUSCO > 98.5%), making them ideal references for *Lupinus* comparative genomics, synteny mapping, and studies of gene family evolution, adaptation, and crop improvement.

Synteny-based Ks analysis revealed a prominent peak at Ks = 0.17 in all six *Lupinus* species but absent in two outgroup legumes, indicating a genus-specific WGD predating *Lupinus* diversification. Gene family dynamics show contracted families outnumber expanded ones in all species, confirming widespread post-WGD diploidization. However, the proportion of WGD-derived genes retained varies strikingly: from 60% in *L. digitatus* to only 36% in *L. mutabilis*, demonstrating distinct lineage-specific trajectories despite a shared WGD.

A core set of WGD-derived functions—actin cytoskeleton organization, ion transport, calcium signaling, and defense responses—is enriched in expanded families across all six species, consistent with the gene balance hypothesis (dosage constraints on macromolecular complexes). Beyond this core, each species exhibits unique signatures: cell wall modification (*L. cosentinii, L. digitatus*), nitrogen metabolism (*L. albus, L. angustifolius*), flower development (*L. luteus*). Strikingly, *L. mutabilis* has lost nearly two-thirds of WGD-derived duplicates (36% retention) and completely lacks the universal cytoskeleton/ion transport enrichments; instead, its expanded families are enriched for MAPK cascade, fatty acid/vitamin E biosynthesis, oxidative stress, and ERAD pathway. We propose this extreme divergence reflects adaptation to its high-altitude Andean habitat (intense UV, temperature fluctuations), favoring a streamlined genome while prioritizing stress signaling and lipid metabolism. Thus, the shared WGD has given rise to both conserved core functions and remarkable lineage-specific adaptive divergence.

## Supplementary Materials

Supplementary Table S1. Statistics of genome assemblies foe the two *Lupinus* species

Supplementary Table S2. BUSCO assessments of the two reassembled and six public genomes

Supplementary Table S3. Statistics of genome annotations

Supplementary Table S4. Infomations of repeat annotations in the genome of *L. cosentinii*

Supplementary Table S4. Infomations of repeat annotations in the genome of *L. digitatus*

Supplementary Table S6. Infomations of WGD, gene family turnover, and WGD-derived genes in expansion families

Supplementary Table S7. Go enrichments of WGD-derived genes in expansion families in *L. cosentinii*

Supplementary Table S8. Go enrichments of WGD-derived genes in expansion families in *L. digitatus*

Supplementary Table S9. Go enrichments of WGD-derived genes in expansion families in *L. albus*

Supplementary Table S10. Go enrichments of WGD-derived genes in expansion families in *L. angustifolius*

Supplementary Table S11. Go enrichments of WGD-derived genes in expansion families in *L. Luteus*

Supplementary Table S12. Go enrichments of WGD-derived genes in expansion families in *L. mutabilis*

## Data Availability Statement

The assembled genome sequences and annotations of *Lupinus cosentinii* and *L. digitatus* are available on figshare at https://doi.org/10.6084/m9.figshare.32032728.

